# Computational analysis of morphological changes in *Lactiplantibacillus plantarum* under acidic stress

**DOI:** 10.1101/2024.11.14.623583

**Authors:** Athira Venugopal, Doron Steinberg, Ora Moyal, Shira Yonnasi, Noga Glaicher, Eliraz Gitelman, Moshe Shemesh, Moshe Amitay

**Affiliations:** Biofilm Research Laboratory, Institute of Biomedical and Oral Research (IBOR), Faculty of Dental Medicine, The Hebrew University of Jerusalem, Jerusalem 9112102, Israel; Department of Food Quality and Safety, Institute for Postharvest Technology and Food Sciences, Agricultural Research Organization, Volcani Center, Rishon LeZion, Israel; Department of Bioinformatic, Jerusalem College of Technology, Jerusalem 9372115, Israel

**Keywords:** *Lactiplantibacillus plantarum*, morphology, deep learning, cell length, object detection, image classification

## Abstract

Cell shape and size often define characteristics of individual or communities of microorganisms in changing environments. Hence, characterizing cell morphology using computational image analysis can aid in the accurate identification of bacterial responses to these changes. Modifications in cell morphology of *Lactiplantibacillus plantarum* were determined in response to acidic stress, specifically during growth stage of the cells at pH 3.5 compared to pH 6.5.

Consequently, we developed a computational method to sort, detect, analyze, and measure bacterial size in a single-species culture. We applied a deep learning methodology composed of object detection followed by image classification to measure the bacterial cell dimensions of the pre-identified cells.

The results of our computational analysis show a significant change in cell morphology in response to alteration of environmental pH. Specifically, we found that the cell was dramatically elongated at low pH, while the width was not altered. Those changes could be attributed to modifications in membrane properties, for instance increased cell membrane fluidity in acidic pH.

Integration of deep learning with microbial microscopic imaging is an advanced methodology for studying cellular structures. These trained models and scripts can be applied to other microbes and cells and are publicly available at: https://github.com/OraMoyal26/bacteria_dimensions/tree/main

## Introduction

The shape of a microorganism is an ecologically important factor because it is associated with many of its properties. Bacteria have a unique capability to maintain precise cell morphology. These morphological characteristics are fundamental metric distinctions between bacteria of different species (1). The morphology of the bacteria is sometimes taken as a constant value. However, variations in bacterial dimensions and shape maybe indicative of the changes in philological, virulence or due to environmental factors (2). In addition to this, morphological modifications may occur with growth rate, stress as changes in temperature values. Apparently, the energy consumed by bacteria also depends on their size (3).

Under certain settings, such as biofilm growth conditions, low carbon bioavailability, carbon-nitrogen imbalance, some bacteria are elongated while maintaining constant width. By elongation, biofilm bacteria strategically enlarge their nutrient collection surface without substantially changing the ratio of surface area to volume (SA/V). This shows that bacteria growing under restricted conditions may morphologically adapt to starvation by elongating (4).

The morphological traits of bacteria are controlled by diverse biochemical pathways and bacterial properties, such as cytoskeletal proteins, elasticity, fluidity of membranes, and osmotic pressure (5). The main common category of bacteria based on their shape is cocci, which may either remain single or attach to one another in groups (streptococci). It may be assumed that coccoid forms were derived from rod-shaped organisms through evolutionary time. Bacilli, which are rod-shaped cells similar to cocci, remain either single or attached to other cells. Another type is the group which includes bacteria that are either helical-shaped or curved (comma-shaped) which can range from slightly curved to corkscrew-like spiral (6)

*Lactiplantibacillus plantarum* is a gram-positive, rod shape bacteria and they thrive in broad arrays of habitat due to their immense ecological and metabolic adaptability (7,8). In addition, *L. plantarum* has gained the status of GRAS (Generally Regarded as Safe) for their valuable application in food and fermentation industry and as potential probiotics and postbiotics organisms (9). The viability and stability of these bacteria can be affected by exposure to stressors such as thermal stress, cold storage, pH imbalance, osmotic shift, and other unstable conditions (10).

*L. plantarum* have adopted different mechanism to tolerate environmental stressors as changes in pH values by; morphological alterations, metabolic changes, growth pattern and genetic modifications (11). Morphological changes in response to stress are considered indicators of survival strategies in bacteria (12). They do so by altering either their cell division pattern or the composition of the peptidoglycan membrane (13,14). Morphological changes in *L. plantarum* can potentially affect its growth kinetics, viability, and adhesion abilities. Furthermore, these morphological alterations can also be advantageous by providing cross-protection against additional environmental changes (15).

As a common response to stress, many rod-shaped bacteria undergo filamentation, wherein the cell division process is hindered, leading to the chaining of the cells without separating from each other (12). However, this type of adaptive morphotype is often overlooked by microscopic observations and lacks precise computational analysis. *L. plantarum* is often exposed to acidic environment as they release lactic acid into the medium during their growth*. L. plantarum* in acidic pH led to the display of phenotypic heterogeneity among the population which enables them to improve their viability (15). A V-shape structure of two bacteria is formed under acid stress (16,17). This morphological change is associated with biofilm formation and is quorum sensing depended via the LUXS / AI-2 pathway (17). We also showed that incomplete cell division process leads to V-shaped multicellular structuring (17). However, detailed quantitative characterization of the morphological changes associated with adaptation to acidic stress was not studied. Hence, computerizing the ultrastructural changes adopted by *L. plantarum* during acid stress will be helpful for exploiting the benefits of this bacterium.

Bacterial morphology is an important microbial parameter that can provide vital information about the properties and ecological stages of bacteria. Several methods have been employed to measure bacterial size. Examples include electron microscopy, coulter counter, flow cytometry epifluorescence microscopy, and transmission electron microscopy. Each of these methods has advantages and disadvantages.

The integration of deep learning into biological microscopy imaging has ushered in a new era of precision and efficiency in the study of cellular structures and processes (18). With the advent of advanced neural network architectures, particularly convolutional neural networks (CNNs), deep learning algorithms have proven invaluable for the automated analysis and interpretation of microscopic images. This integration allows characterizations such as cell (or subcellular organelle) classification, segmentation, and detection to be performed with unprecedented accuracy and speed (19). Size estimation involves the process of determining the dimensions or physical size of objects within an image. This task is fundamental in various applications ranging from fruit size (20,21) to cell dimension analysis (22,23).

One of the more advanced means to measure bacterial cells dimensions is by deep learning algorithms using computational vision. In this study, we used two computer vision methods: image classification and object detection. Image classification categorizes images into predefined classes based on their visual content. Object detection involves the identification and localization of objects within images. The goal is not only to recognize the types of objects present in an image but also to provide precise bounding box coordinates around each detected object. Both methods consist of training on datasets with labeled images to allow the algorithm parameters to capture intricate patterns and variations, allowing them to generalize well to new, unseen images.

The aim of this study is to quantitatively characterize the changes in *L. plantarum* morphology (16,17) by computational analysis. Our results provide a novel computational approach for detecting differences in morphological changes adopted by *L. plantarum* during adaptation to acidic stress.

## Materials and methods

### Bacterial strains and growth conditions

*L. plantarum* 12422 (Lallemand, Blagnac, France) was used in this study for analysis and morphological characterization. For routine growth, *L. plantarum* was cultured in De Man, Rogosa, and Sharpe (MRS) medium (HiMedia Laboratories Pvt. Ltd., Maharashtra, India) at pH 6.5, or on MRS agar at 37 °C with 5% CO_2_, under non-shaking conditions. For acidic pH stress conditions, the MRS medium was adjusted to pH 3.5 using 1M HCl. For each experiment, an overnight culture of *L. plantarum* cultured at pH 6.5 (OD_600_ approx. 2) was diluted to an OD_600_ of 0.1 using MRS medium of either pH 6.5 or 3.5.

### High-Resolution Scanning Electron Microscopy (HR-SEM)

HR-SEM (Electron Microscopy Sciences, Hatfield, PA) was used to image *L. plantarum*, when cultured in MRS-3.5 and MRS-6.5. The bacterial cells were collected after 24 h of growth, centrifuged (5000 × g, 5 min, 4 °C), fixed in 4% glutaraldehyde (Electron Microscopy Sciences, Hatfield, PA, USA) in sterile water for 2 h, and washed twice with sterile water before letting them dry on a 0.5 × 0.5 cm glass slide. The samples were coated with iridium prior to imaging using a Cryo High-Resolution Scanning Electron Microscope (Apreo 2S, Thermo Fischer Scientific) at 5000X and 20,000X magnification. All samples were tested in biological and experimental triplicate.

### Cell dimensions analysis from SEM images

To measure bacterial cell dimensions, we utilized deep learning techniques from the field of computer vision. Several challenges were encountered in the dimension measurement of noisy rod-shaped cell images (Fig. 1).

1. Incomplete cells because of overlapping cells or partial cells near the borders of the image.
2. Cells during mitosis (e.g., actually two cells, not a single cell).

**Figure 1.**
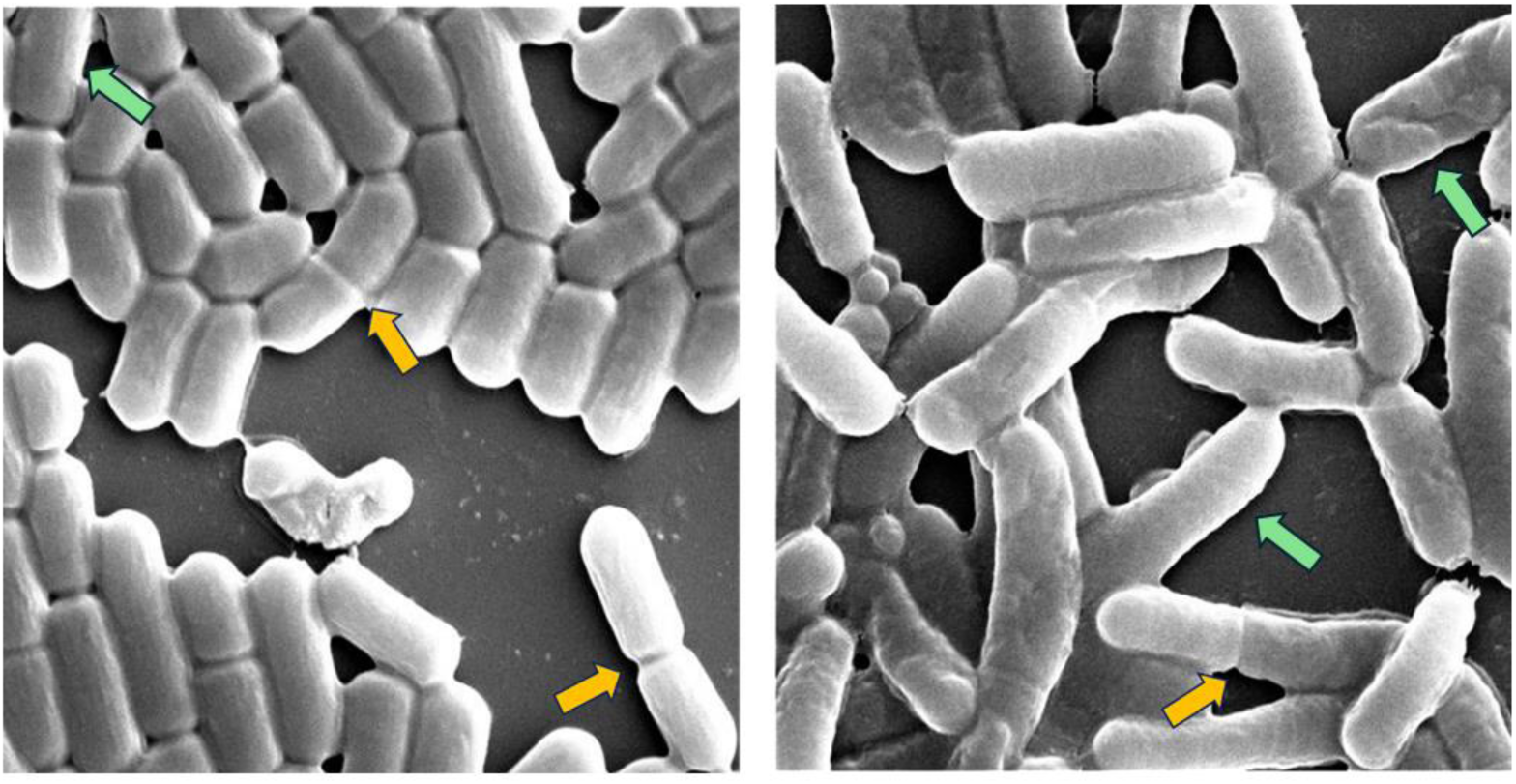
Examples of cells during mitosis (yellow arrows) and incomplete divided cells (green arrows). Left: control; right: acidic condition.

We did not use a specialized analysis software (such as <u>SuperSegger</u>) since we wanted to make sure that our methodology properly addressed these challenges (24). The following methodology was applied to measure bacterial cell dimensions (depicted in Fig. 2). We used object detection (identification and localization of objects within images) to identify and locate bacterial cells in the SEM images (25). We then applied image classification (training of an algorithm to recognize patterns in images that distinguish one class from another) to select only the bacterial bounding boxes whose dimensions represent the dimensions of the bacterial cells. At this stage, we filtered out partial cells and cells during mitosis. In the following step, we applied the Canny algorithm to detect the edge of the bacterial cell and calculate the width and length of the cell. We evaluated the computational results by using manual measurements that were not included in the training. We used this methodology to compare the width and length of bacterial cells under the two conditions.

**Figure 2.**
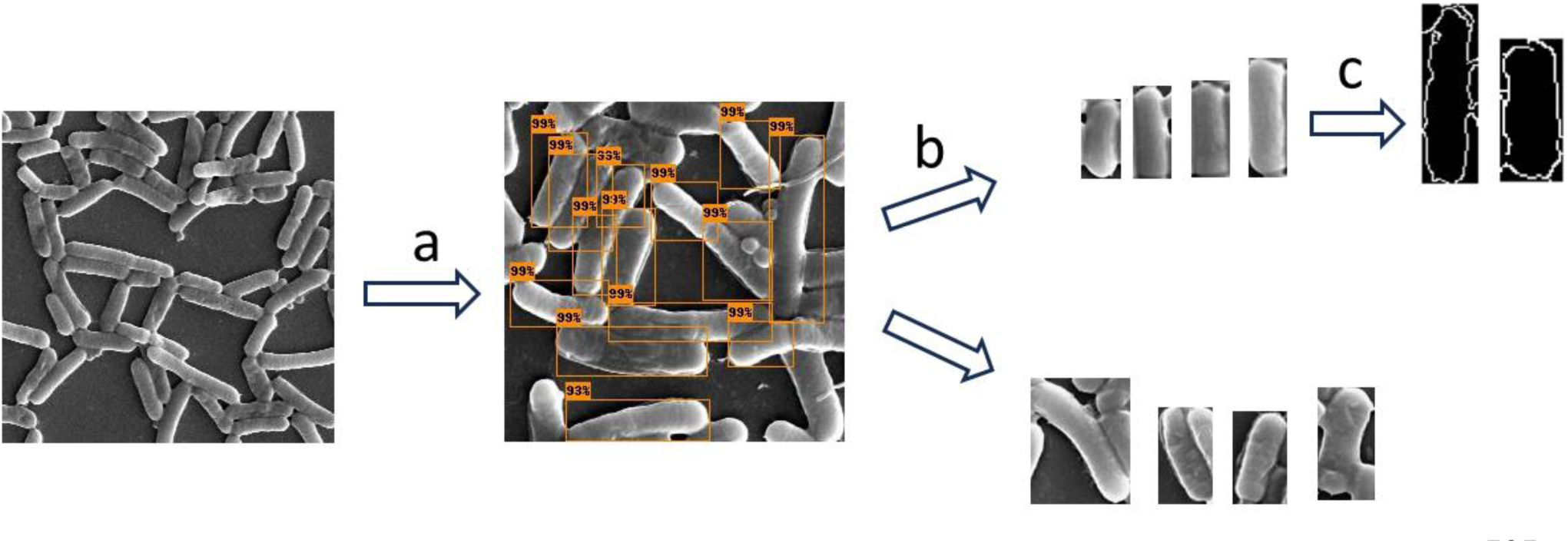
A diagram illustrating the workflow undertaken to measure the bacterial cell dimensions: First, (a) object detection was applied to isolate the cells from the SEM image. Step (b) Image classification to pick cell images with image dimensions representing the dimension of the bacteria, and final step (c). Canny edge detection algorithm was applied to obtain more accurate measurements.

### Image classification: acidic vs. control

SEM images were obtained at 5 K magnification. The initial SEM slides were modified from 1500 × 1000 pixels to 24 images of 250 × 250 pixels. 20, 200 and 4,400 magnitude images were used for training section, validation and test respectively. The image classification model was implemented using Keras version 2.15.0 (26). Fine tuning was performed with the pretrained EfficientNetB0 architecture (27). The following top layers were added: GlobalAveragePooling2D, a dense layer with 16 neurons, dropout of 0.15, and an output layer of two neurons (28). The images were pre-processed by resizing to a uniform size of 128X128 pixels and normalizing pixel values. Data augmentation techniques such as vertical and horizontal flipping have been applied. The model was trained (only the top layers) using an Adam optimizer (29) with a polynomial decay schedule of the learning rate, using an initial learning rate of 10^-3^. The categorical cross-entropy loss function was chosen as the optimization criterion.

### Measurement of bacterial dimensions

#### Step A: Object detection

Object detection was conducted using images from 20 K SEM magnification. The image size is 500 × 500 pixels. We manually annotated the bounding boxes of only full-length bacterial cells (a single class). The dataset consisted of 135 images from both the acidic and control conditions (an equal quantity from each condition). The dataset was split into training validation and test sets with quantities of 50, 35, and 50. Data augmentation was performed by applying horizontal and vertical flips, zooming and adjusting brightness and contrast.

Faster RCNN (30) architecture was applied with Resnet-50 as backbone and Feature Pyramid Networks (FPN) of TensorFlow Model Garden (TFM) module of TensorFlow 2. The performance of the trained model on the test set was: average precision, mAP (0.5-0.95), of 0.71 and AP_50_ of 0.88 (31). This object detection model was then used to obtain more than 25 K images of separate bacterial cells for the next step of image classification.

### Step B - Image classification

Image classification was conducted using images from the object detection output bounding boxes. We manually created two classes: 1. Cell images that their image dimensions represent their cell dimensions and 2. Cell images that their image dimensions did not represent their cell dimensions. (Sample images depicted in Fig. 2, Step 3) cells of both acidic and control conditions were included (an equal quantity from each condition). The dataset was randomly split into train, validation and test sets. Each set consisted of 840, 240 and 120 respectively and had an equal quantity from each class.

The image classification model was implemented using Keras version 2.15.0 (26). Fine tuning of the pretrained EfficientNetV2 (27) architecture was performed. The following top layers were added: GlobalAveragePooling2D, dense layer with 16 neurons, dropout of 0.15 and an output layer of 2 neurons. The images were preprocessed by resizing to a uniform size of 128X128 pixels and normalizing pixel values. We applied data augmentation techniques such as vertical and horizontal flipping.

The model was trained (only the top layers) using an Adam optimizer (32) with polynomial decay schedule of learning rate, using an initial learning rate of 10^-3^. The training process involved 30 epochs, and the batch size was set to 16. The categorical cross-entropy loss function was chosen as the optimization criterion. The final accuracy of the trained model on the test set was 0.88. We then used this classifier to filter out from more than 25 K single cell images the images that their image dimensions represented their cell dimensions and got 1200 images. A manual examination was performed on these images and created a final dataset of 600 images (i.e 300 images from each condition: acidic and control). Additionally, Canny edge detection algorithm (33) was applied to get a more accurate measurement of the width (using OpenCV Python package version 4.10) (27). In order to further evaluate our results, we manually measured the length and width of 200 cells from this dataset.

All the code and the models are available in the following repository: https://www.computer.org/csdl/journal/tp/2017/06/07485869/13rRUx0gera.

### Growth kinetic studies

Bacterial growth profile of *L. plantarum* in MRS-6.5 and MRS-3.5 was evaluated by kinetic studies. An overnight culture of *L. plantarum* was diluted in 10 ml of either MRS-6.5 or MRS-3.5 medium to an initial OD_600nm_ of 0.1. 200 µl of this culture were inoculated in a well sterile tissue grade transparent flat-bottomed 96-well microplate (Corning, Incorporated, Kennebunk, ME, USA) and bacterial growth was measured at an OD_600_ nm at every 30 min for a period of 24 h in a Tecan M200 infinite microplate reader (Tecan trading AG, Mannesdorf, Switzerland). The experiment was performed in triplicates and expressed as mean ± standard deviation.

### Laurdan membrane fluidity assay

Changes in membrane fluidity associated with morphological changes was recorded using a fluorescent probe Laurdan, which intercalates with the peptidoglycan bilipid layer and emits a shift in the wavelength depending on the number of water molecules (34). 1 ml of OD 0.3 of L. *plantarum* suspension, cultured in pH 6.5 and pH 3.5 for 4 hours, were centrifuged. Then, Laurdan (AnaSpec Inc., Fremont, CA, USA) was added to a final concentration of 10 µM, and the samples were incubated for 10 min at 37°C. As controls, unstained samples of pH 6.5 and pH 3.5 were also prepared. After incubation, the bacteria were washed four times with a mix of 1 mL PBS and 0.1% dimethylsulfoxide (DMSO). The fluorescence emitted from 200 µL of each sample, in triplicates, in a µ-clear black flat 96-well plate (Greiner Bio-One) was monitored in the M200 infinite plate reader (Tecan, Trading AG, Männedorf, Switzerland) with an excitation of 350 nm and emission spectrum from 400 nm to 600 nm at 30 °C. Membrane fluidity was measured using Laurdan generalized polarization (GP) values according to the formula GP= (RFI_440nm_ − RFI_490nm_)/ (RFI_440nm_ + RFI_490nm_) as described (34).

## Results

### 1. Image classification method shows that *L. plantarum* shows significant differences in morphology when cultured in acid stress

With the aim to examine the morphological changes in acidic condition compared to the control, we performed SEM imaging of *L. plantarum* and applied image classification method. We trained the algorithm to learn features and patterns in the images to accurately classify the two environmental states of control and acidic conditions (Fig 3 presents sample images from each state). To demonstrate that the two environmental conditions are significantly different, we used 20 images from the SEM for training (10 of each state). Additionally, the training images were taken from a single cultivation batch **(Fig 3B)** and tested on the two other cultivation batches.

**Figure 3.**
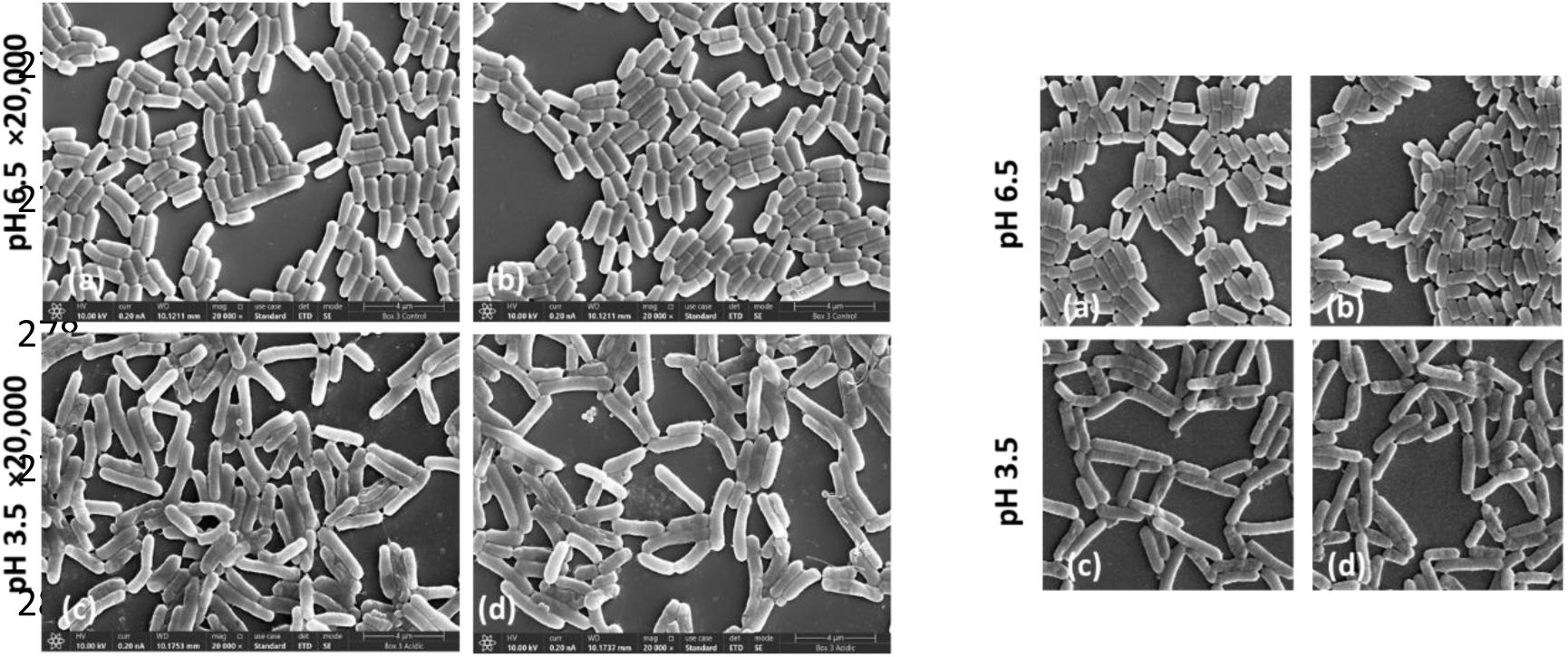
Sample images of the control (pH 6.5) and acidic (pH 3.5) used for training the algorithm.

Our results demonstrated a near perfect prediction accuracy (number of correct predictions out of total number of predictions) of the unseen test set of 0.97 (The detailed confusion matrix is available in Table 1). The results showed that 97% of the 250X250 pixels images that were cut from the 5K SEM magnification images in the acidic condition were different from the control. Bearing in mind that the training was done on only 20 images, we can conclude that the morphology differences between the two conditions are consistent and significant.

**Table 1.** Confusion matrix of the image classification classifier – control Vs. acidic.

| Test samples | Predicted control | Predicted acidic |
| --- | --- | --- |
| Actual control | 2182 | 18 |
| Actual acidic | 116 | 2084 |

### 2. Acidic stress leads to slower growth and increase in length of *L. plantarum*

Next, we examined whether the morphological changes between the acidic and control samples had any relation with the growth of the *L. plantarum* cells in those pH conditions. Exposure of *L. plantarum* to pH 6.5 and pH 3.5 showed significant differences in their growth pattern.

Additionally, manual measurements of the length and width from the SEM images of *L. plantarum* at pH 6.5 and pH 3.5 (**Fig 4.C**) shows that the length of the cells at pH 3.5 has comparatively increased after 24 h of growth (Fig 4C (c-d)). However, at pH 6.5, the cells remained a single, rod-shape cells, with no increase in length (**Fig 4C. (a-b)**). As shown before (17), manual recording of the planktonic growth of *L. plantarum* in 50 ml Eppendorf tube at pH 3.5 showed a longer lag phase and after 24 h of growth, the cells in pH 3.5 the continued to grow exponentially (17). We observed a similar trend in the growth pattern when cultured in a 96-well plate **(Fig 4B).** Cell elongation together with the increase in the growth in pH 3.5 suggests that *L. plantarum* undergoes an adaptational change in acid stress.

**Figure 4:**
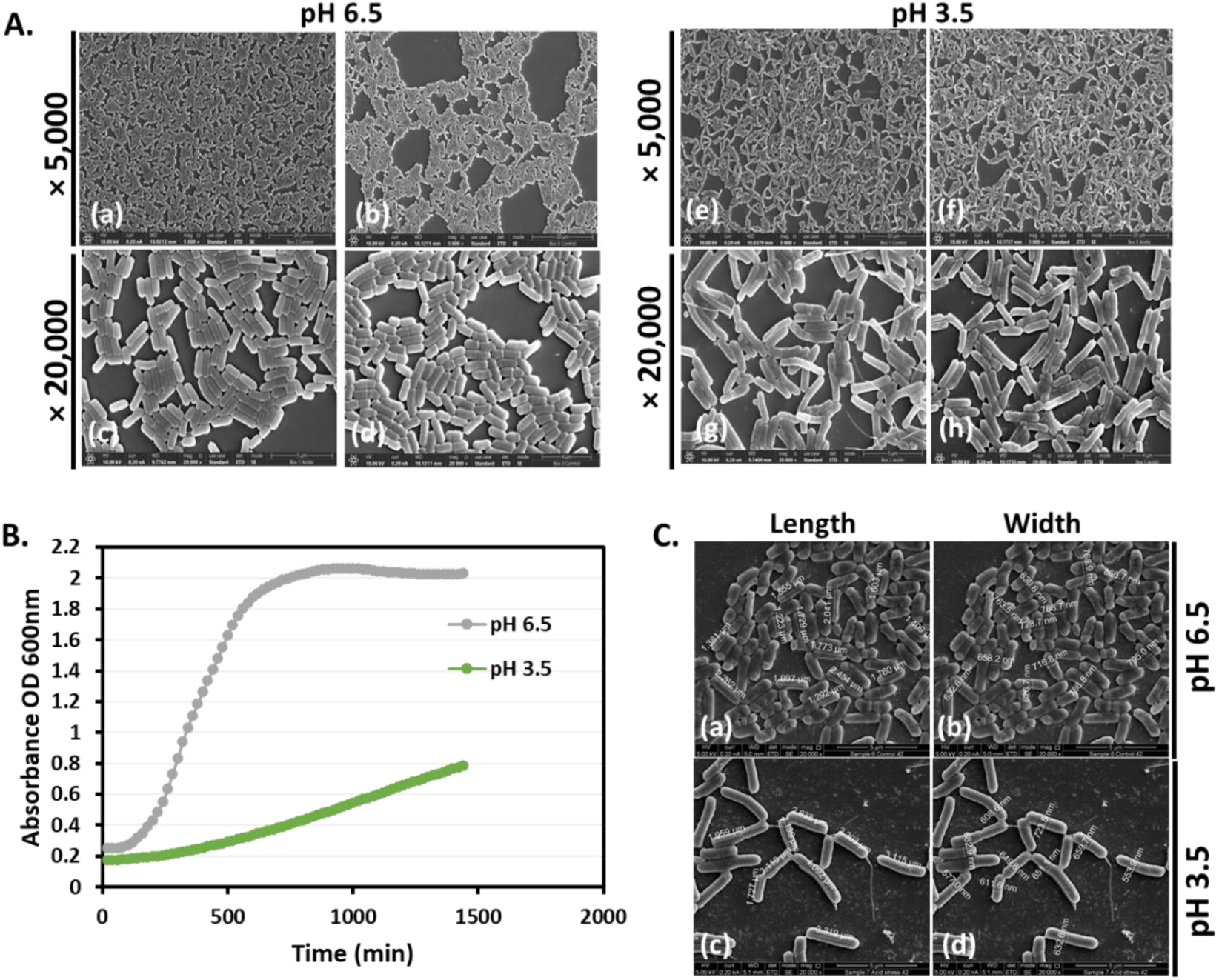
*L. plantarum* adapts to acidic stress by increasing their cell length. A. SEM images of *L. plantarum* at pH 6.5 at magnifications (a-b) ×5000 (c-d) ×20,000 and pH 3.5 at magnifications (e-f) ×5000 (g-h) ×20,000 after 24 h of incubation at 37°C. B. Growth kinetics of *L. plantarum* in pH 6.5 and pH 3.5 for a period of 24 h on a 96-well tissue grade polystyrene plate. C. Manual measurements of length and width of *L. plantarum* at pH 6.5 (a-b) and pH 3.5 (c-d) on SEM images.

### 3. Comparison of cells dimensions in control Vs. acidic conditions

#### Bacterial dimensions analysis

We compared the length and width dimensions of the bacteria’s cells at the acidic and control environments. The accuracy of our computational approach was evaluated by comparing its measurements to the manual measurements. We obtained reasonable root mean square error (RMSE) deviations of 2.66 pixels for the length and 1.7 pixels for the width. This indicates a consistency between the computational and the manual measurements.

Table 2 shows the cells dimensions based on manual measurements comprised of 200 cells (100 cells from each experimental group). The median length of the acidic bacteria increased by 39% whereas the width of the bacteria did not undergo significant change. Moreover, the variation of the length nearly doubled at the acidic conditions.

**Table 2.**
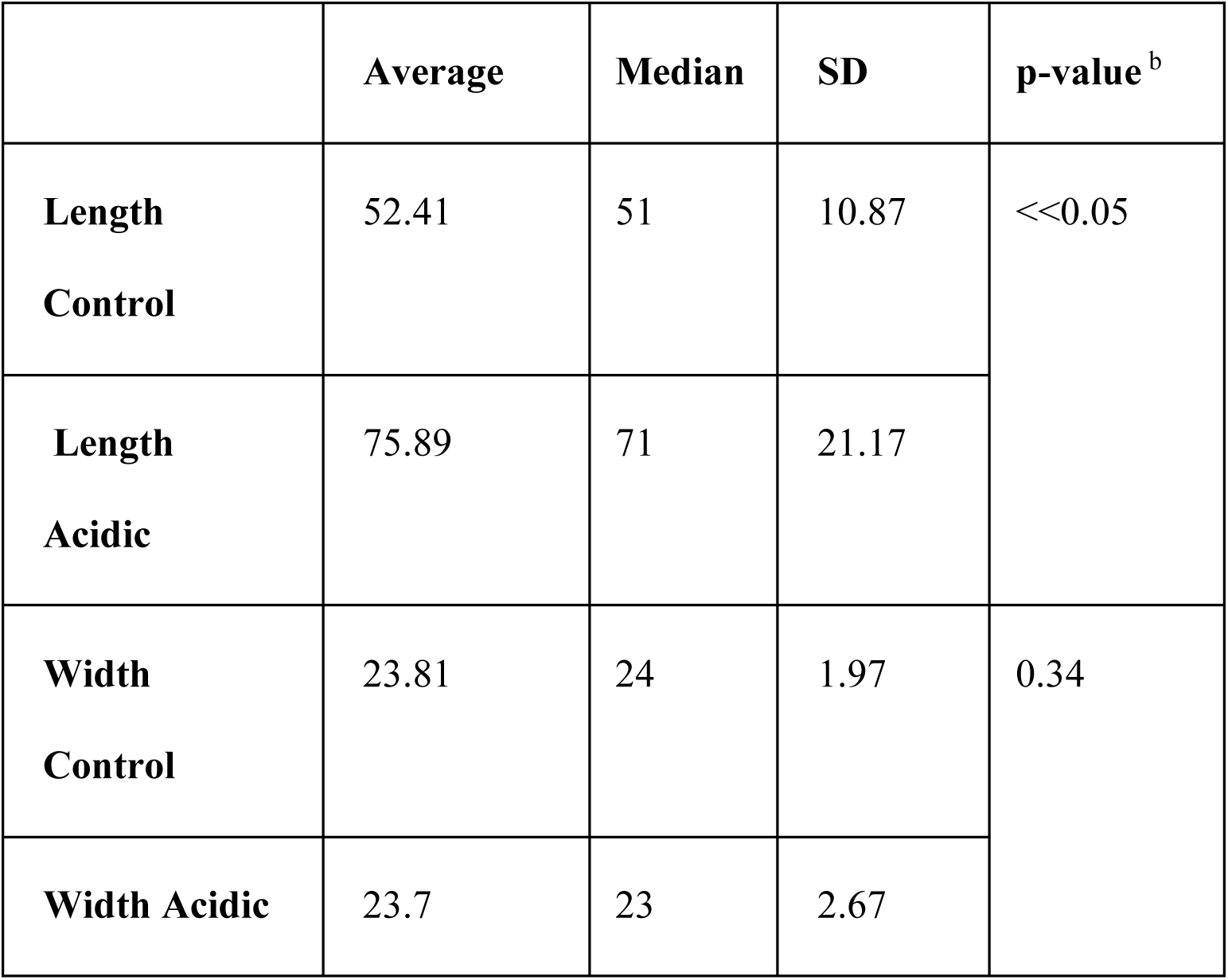
Comparison of cells dimensions in control Vs. acidic conditions - *manual measurements*. ^a^.

**Table 3.**
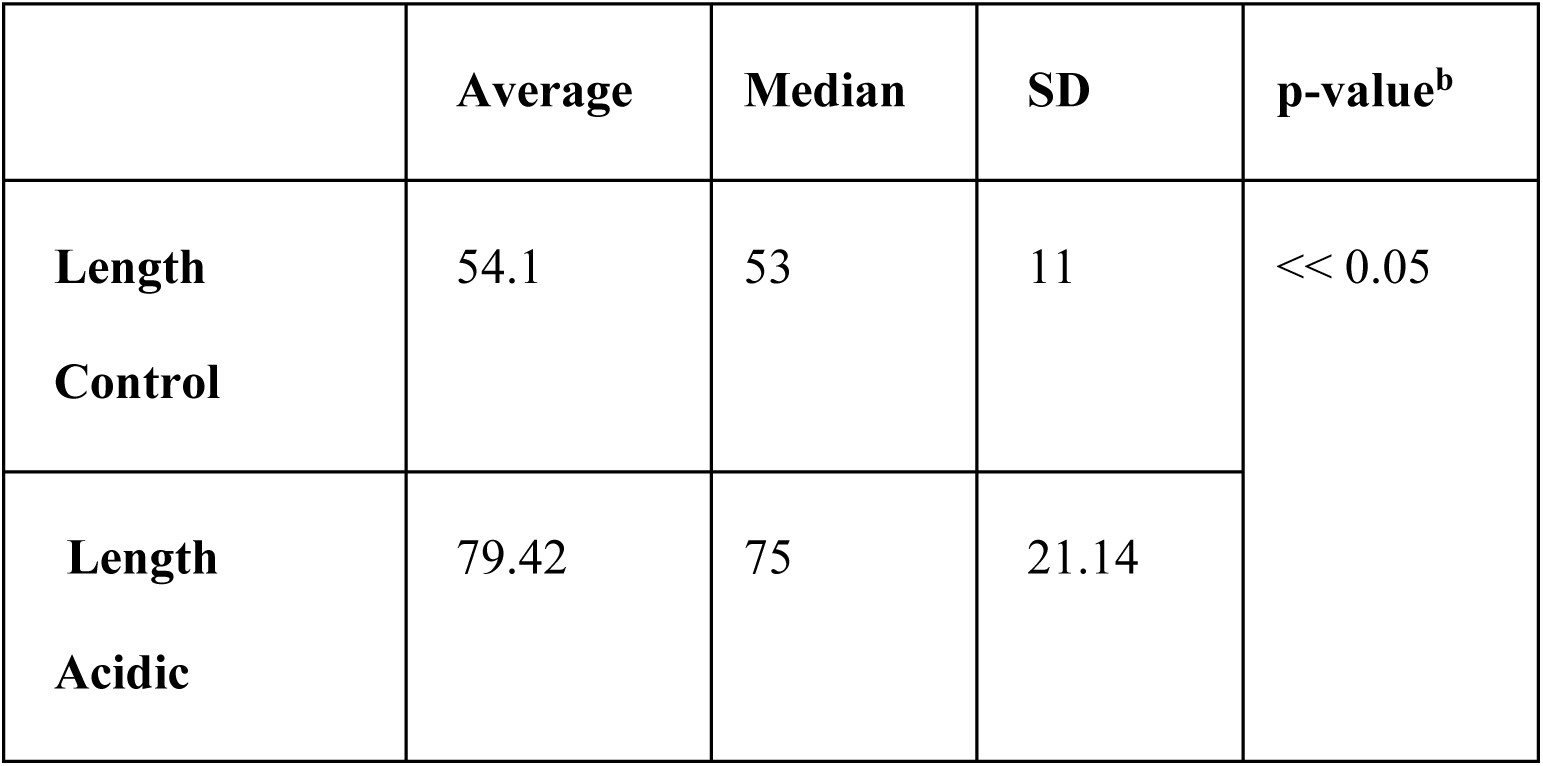
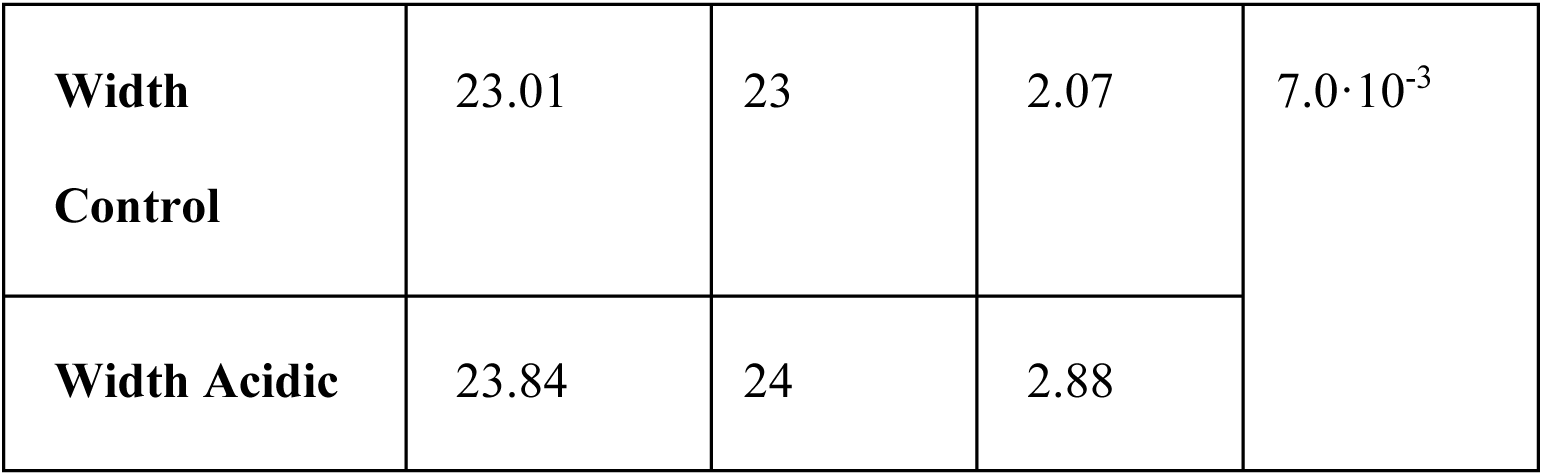
Comparison of cell dimensions in control Vs. acidic conditions *–* computational measurements.^a^.

Table 2 and Figure 5 show a summary of the cell’s dimensions in the acidic and control conditions based on computational methodology comprised of 600 cells (300 cells from each experimental group). The computational results were in agreement with those of the manual measurements (table 2): the length of the median acidic bacteria increased by 41% whereas the width of the bacteria underwent a significant change of only a single pixel. The larger sample size in the computational measurements increased the sensitivity to detect small effects, however a single pixel difference has no biological meaning. Additionally, the variation of the length was shown to nearly doubled at the acidic conditions.

**Figure 5.**
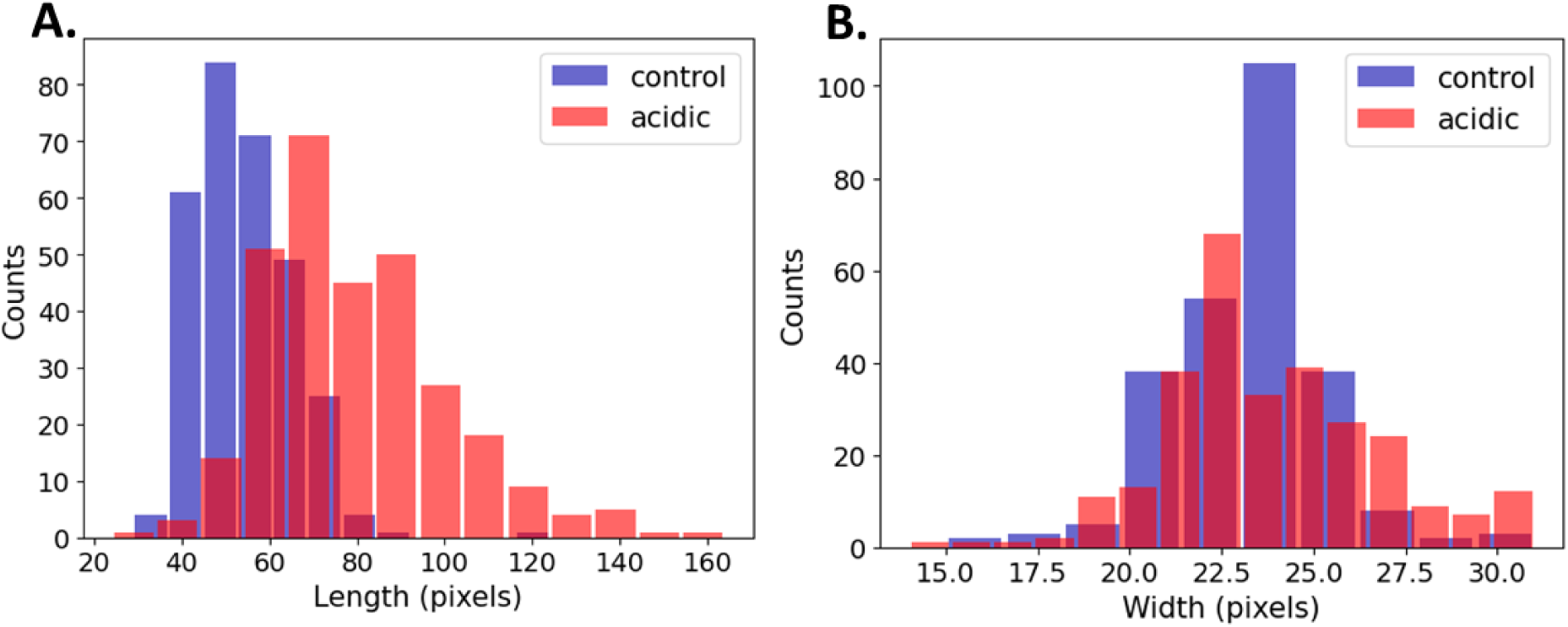
Histograms of length (Left) and width (right) of control Vs. acidic conditions.

### 4. Membrane fluidity in *L. plantarum* is enhanced by acidic stress

As it is apparent that a change in cell morphology occurs, we tested the fluidity of the membrane in the different pH conditions. By visual observation, we can observe that at pH 3.5 the cells share their outer membrane with the neighboring cells with no clear boundaries at the end of the cells (**Fig 6.D marked in red**). However, at pH 6.5, the membrane between two cells can be clearly distinguished from each other (**Fig 6.C (a-b)).** To further substantiate the finding, we performed the bacterial membrane fluidity assay by harvesting the bacteria after 24 h from pH 6.5 and pH 3.5 and staining them with the Laurdan stain, which is a fluorescent probe that intercalates to the region in the peptidoglycan membrane and displays an emission wavelength shift depending on the number of water molecules present between them. The relative fluorescence intensity which measures the degree of membrane fluidity is inversely proportional to the generalized polarized value (GP). Further, increase in Laurdan staining indicates increased membrane fluidity. Our results show that at pH 3.5 the cells are stained more with the Laurdan stain compared to that of pH 6.5 indicating that there is an increase in membrane fluidity at pH 3.5 (**Fig 6. A**). Also, the generalized polarized (GP) values (**Fig 6. B**) shows that at pH 3.5 the cells have lesser GP value supporting the result of increased membrane fluidity.

**Figure 6:**
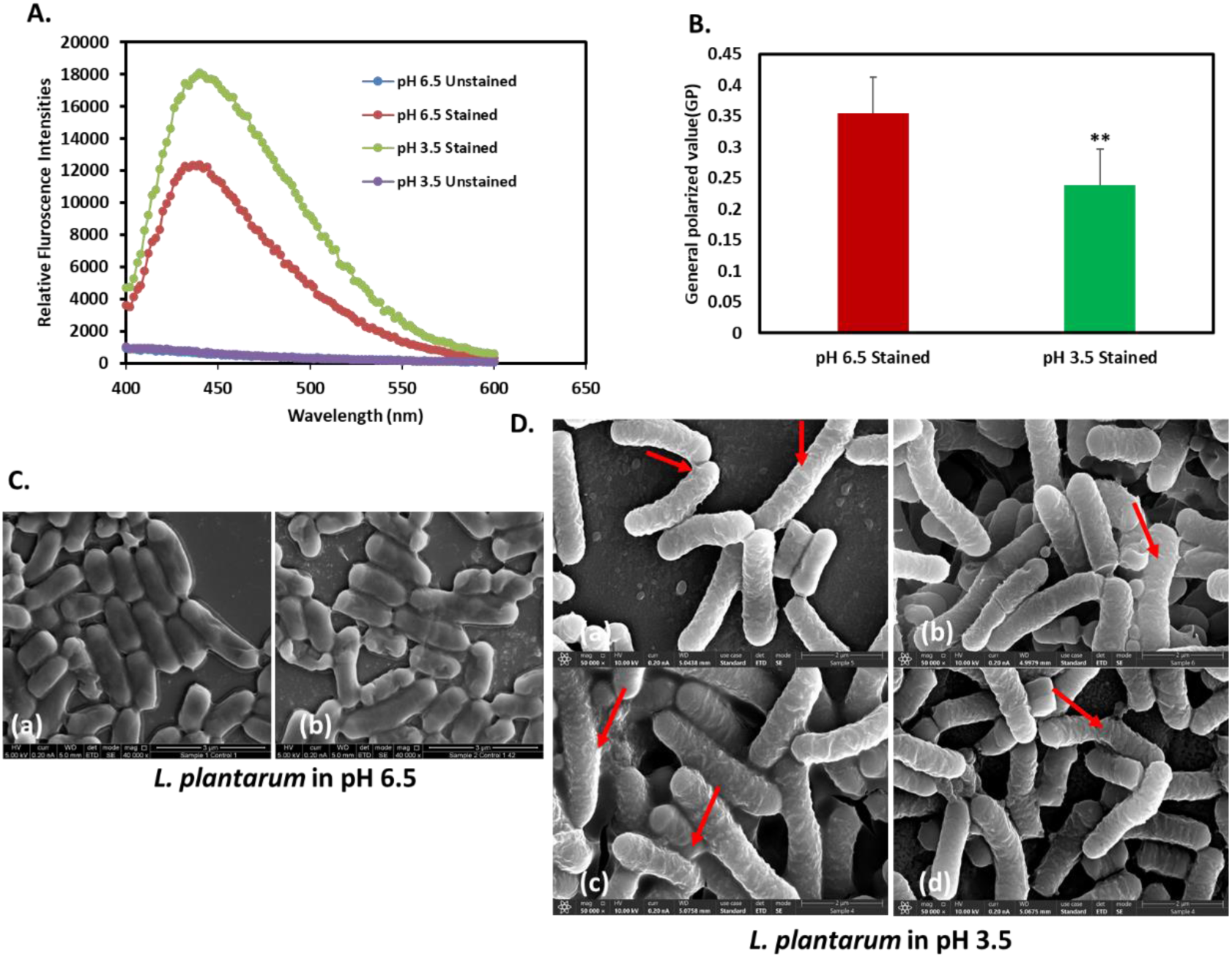
Acid stress induces changes in membrane fluidity in *L. plantarum*. Membrane fluidity assay expressed as: A. relative fluorescence intensities and B. generalized polarized (GP) value C. High-resolution SEM images of L. plantarum cells at pH 3.5 at × 50,000 magnification. Red arrows indicate rough membrane surface (a-b) and fluid membrane (c-d).

## Discussion

Bacterial morphology is a fundamental criterion for microbial classification. Variations in cell morphology under diverse environmental conditions may signal alterations in bacterial properties. As our understanding of bacterial ecological niches deepens, it is increasingly evident that bacteria can modify their morphology in response to environmental fluctuations. Certain cell morphologies are now recognized as advantageous for survival in dynamic or hostile micro-environments, which is particularly significant in the context of host colonization and infection.

Numerous bacterial pathogens employ morphological changes as a strategy to evade host immune responses and persist in specific host sites or adapt to environmental changes. For instance, the size of *Endomyces magnusii* has been observed to vary in response to different media conditions. *Endomyces magnusii* size was altered in the presence of different media. Cells exposed to 2% glucose were 30-35% larger than those cells growing on glycerol. However, a decrease of the initial glucose concentration to 0.5-0.2% only slightly altered cell length (35). Enterobacter sp. strain, SM1_HS2B, growing in the Luria-Bertani medium, delay their cell division and present elongate morphology compare to standard growth conditions (36). Bacillus sp. SFC 500-1E has developed adaptive strategies as an increase in bacterial size, such as length, width, and height, in the presence of Cr(VI) + phenol treatment. In addition, its biofilm was also modified. Rapid rigidity of the cell membrane, was induced in Cr(VI) + phenol treatment. This effect was counteracted after 16 h of incubation (37). We have also shown that changes in environmental pH have resulted in a morphological change of *Lactiplantibacillus plantarum* with the cells mainly changing to a V-shaped structure of 4-8 cells attached to a common point (16,17). Apart from changes in the membrane fluidity and morphological changes we observed a slow increase in growth of *L. plantarum* when cultured in pH 3.5 on a 96-well microplate (Corning, Incorporated, Kennebunk, ME, USA). However, after 24 h of growth, the cells continued to grow exponentially when compared to that of cells cultured in pH 6.5. A similar trend in growth profile was also observed previously (17) when cells were cultured in bigger volumes of 50 ml Eppendorf tubes.

In this study, we demonstrated by computational analysis that the morphology of *L. plantarum* differs significantly in an acidic environment compared to control conditions. Remarkably, an image classification model trained with only 20 images achieved near-perfect accuracy in predicting the growth environment depicted in the images. We then focused on three key features: growth rate, membrane fluidity, and cell dimension analysis. Our findings indicate that membrane fluidity increases under acidic pH conditions compared to the control. This increase in fluidity may facilitate morphological changes, such as cell elongation. It is plausible that membrane properties, which are integral to maintaining bacterial shape, undergo modifications during morphological adaptations. Furthermore, the morphology of other bacteria also appears to be sensitive to environmental pH changes. For example, acidic conditions influenced the length at which *Escherichia coli* initiated the division process (38).

In our study, we employed computational analysis to more accurately assess the size differences of *L. plantarum* under acidic stress compared to standard growth conditions. In this study we sampled a large population of bacteria that we obtained from the SEM images and categorized the images as the control and test groups. However, to minimize false positives and negatives, the algorithm was trained to exclude images of bacteria that did not exhibit clear and well-defined geometric features. By using modified algorithms tailored to specifically characterize and measure dimensional features, we obtained more precise information on changes in cell dimensions.

We have developed a computational methodology for measuring the width and length of rod-shaped bacteria, with the primary goal of comparing cell dimensions under two conditions. This methodology involved object detection followed by image classification. By utilizing computerized imaging, we achieved quick, accurate, and reliable detection of cell characteristics within a culture.

We applied this methodology to sample and measure cell dimensions, achieving results closely aligned with manual measurements—showing a size increase of 41% versus 39%, respectively. The image classification effectively filtered out partial and dividing cells. Importantly, the same computational procedure, with identical parameter settings, was applied to each condition, ensuring that any bias or discrepancies were consistently reflected across both conditions. Additionally, this methodology does not require extensive or complex dataset labeling, as both object detection and classification tasks are straightforward, making it relatively easy to implement. Most studies on size determination use object detection to isolate objects within images, with some employing semantic segmentation to define precise boundaries for accurate measurement of size or diameter. In this study, we opted for image classification to specifically address the need to classify and exclude partial, overlapping, and dividing cells. All code and trained models from this study are available in the GitHub repository and can be used for the dimensional analysis of other rod-shaped cells. The methods developed hereby provide another tool to detect and measure bacteria morphology in a cell culture. This method can be applied to other bacteria or cells at different environmental conditions.

## Acknowledgements

We would like to thank Dr. Ronit Vogt Sionov and Dr. Yulia Krouptiski for their scientific support. We are grateful to Dr. Vitaly Gutkin at the Hebrew University Center for Nanoscience and Nanotechnology, for his assistance with the SEM analysis.

## Author Contributions

**Athira Venugopal:** Conceptualizing, Methodology, Investigation, Conducting, Data curation and analysis, Writing-Original draft, Writing-Reviewing and Editing. **Shira Yonnasi:** Conceptualizing, Methodology, Investigation, Formal Data Analysis. **Noga Glaicher:** Conceptualizing, Methodology, Investigation. **Eliraz Gitelman:** Conceptualizing, Methodology, Investigation**. Doron Steinberg:** PI, Conceptualizing, Investigation, Methodology, Formal Data Analysis, Writing-Reviewing and Editing, Supervision, Project administration, Resources, Funding Acquisition. **Moshe Shemesh:** PI, Conceptualizing, Investigation, Methodology, Formal Data Analysis, Writing-Reviewing and Editing, Supervision, Project Administration, Resources, Funding Acquisition. **Moshe Amitay:** PI, Conceptualizing, Investigation, Methodology, Formal Data Analysis, Writing-Reviewing and Editing, Supervision, Project administration, Resources, Funding Acquisition

## Funding

This work was supported by internal budgets of all three PIs.

## Declaration of competing interests

The authors declare that they have no competing interest.

## Notes

### Competing Interest Statement

The authors have declared no competing interest.

## References

1. Justice, S. S., Harrison, A., Becknell, B., & Mason, K. M. (2014). Bacterial differentiation, development, and disease: mechanisms for survival. FEMS microbiology letters, 360(1), 1–8. 10.1111/1574-6968.12602

2. Chrzanowski, T. H., Crotty, R. D., & Hubbard, G. J. (1988). Seasonal variation in cell volume of epilimnetic bacteria. Microbial ecology, 16(2), 155–163. 10.1007/BF02018911

3. Determination of Size and Morphology of Aquatic Bacteria by Automated Image Analysis *ByRoland Psenner* BookHandbook of Methods in Aquatic Microbial Ecology, 1^st^ Edition, 1993, Imprint CRC Press, ISBN9780203752746

4. Steinberger, R. E., Allen, A. R., Hansa, H. G., & Holden, P. A. (2002). Elongation correlates with nutrient deprivation in Pseudomonas aeruginosa-unsaturates biofilms. Microbial ecology, 43(4), 416–423. 10.1007/s00248-001-1063-z

5. Jiang, H., & Sun, S. X. (2010). Morphology, growth, and size limit of bacterial cells. Physical review letters, 105(2), 028101. 10.1103/PhysRevLett.105.028101

6. Young K. D. (2007). Bacterial morphology: why have different shapes? Current opinion in microbiology, 10(6), 596–600. 10.1016/j.mib.2007.09.009

7. Svetoslav Dimitrov Todorov & Bernadette Dora Gombossy De Melo Franco (2010) Lactobacillus Plantarum: Characterization of the Species and Application in Food Production, Food Reviews International, 26:3, 205–229, DOI: 10.1080/87559129.2010.484113

8. Yilmaz, B., Bangar, S. P., Echegaray, N., Suri, S., Tomasevic, I., Manuel Lorenzo, J., Melekoglu, E., Rocha, J. M., & Ozogul, F. (2022). The Impacts of *Lactiplantibacillus plantarum* on the Functional Properties of Fermented Foods: A Review of Current Knowledge. Microorganisms, 10(4), 826.

9. Behera, S. S., Ray, R. C., & Zdolec, N. (2018). Lactobacillus plantarum with Functional Properties: An Approach to Increase Safety and Shelf-Life of Fermented Foods. BioMed research international, 2018, 9361614. 10.1155/2018/9361614

10. Dlangalala, T.N.; Mathipa-Mdakane, M.G.; Thantsha, M.S. (2022) The Morphological and Functional Properties of Lactiplantibacillus plantarum B411 Subjected to Acid, Bile and Heat Multi-Stress Adaptation Process and Subsequent Long-Term Freezing. Microbiol. Res. 2022, 13, 909–927. 10.3390/microbiolres13040064

11. Ricciardi A, Parente E, Guidone A, et al. Genotypic diversity of stress response in Lactobacillus plantarum, Lactobacillus paraplantarum and Lactobacillus pentosus. Int J Food Microbiol. 2012;157(2):278–285. doi:10.1016/j.ijfoodmicro.2012.05.018

12. Karasz DC, Weaver AI, Buckley DH, Wilhelm RC. Conditional filamentation as an adaptive trait of bacteria and its ecological significance in soils. Environ Microbiol. 2022;24(1):1–17. doi:10.1111/1462-2920.15871

13. Deghorain, M., Fontaine, L., David, B., Mainardi, J. L., Courtin, P., Daniel, R., Errington, J., Sorokin, A., Bolotin, A., Chapot-Chartier, M. P., Hallet, B., & Hols, P. (2010). Functional and morphological adaptation to peptidoglycan precursor alteration in Lactococcus lactis. The Journal of biological chemistry, 285(31), 24003–24013. 10.1074/jbc.M110.143636

14. Parlindungan, E., May, B. K., & Jones, O. A. H. (2019). Metabolic Insights Into the Effects of Nutrient Stress on Lactobacillus plantarum B21. Frontiers in molecular biosciences, 6, 75. 10.3389/fmolb.2019.00075

15. Ingham, C. J., Beerthuyzen, M., & van Hylckama Vlieg, J. (2008). Population heterogeneity of Lactobacillus plantarum WCFS1 microcolonies in response to and recovery from acid stress. Applied and environmental microbiology, 74(24), 7750– 7758. 10.1128/AEM.00982-08

16. Rajasekharan, S. K., & Shemesh, M. (2022). Spatiotemporal bio-shielding of bacteria through consolidated geometrical structuring. NPJ biofilms and microbiomes, 8(1), 37. 10.1038/s41522-022-00302-2

17. Venugopal, A., Sionov, R.V. Kroupitski, Y., Steinberg, D., Shemesh, M. (2024). The V-shaped structuring regulated via a LuxS-dependent quorum-sensing pathway is associated with Lactiplantibacillus plantarum survivability in acidic environment. Food Frontiers, 10.1002/fft2.514

18. Zhang, J., Li, C., Rahaman, M.M. et al. (2022) A comprehensive review of image analysis methods for microorganism counting: from classical image processing to deep learning approaches. Artif Intell Rev 55, 2875–2944. 10.1007/s10462-021-10082-4

19. Litjens G, Kooi T, Bejnordi BE, Setio AAA, Ciompi F, Ghafoorian M, van der Laak JAWM, van Ginneken B, Sánchez CI. (2017) A survey on deep learning in medical image analysis. Med Image Anal. Dec; 42:60–88. doi: 10.1016/j.media.2017.07.005. Epub 2017 Jul 26. PMID: 28778026.

20. Ma, P., Li, C., Rahaman, M.M. et al. A state-of-the-art survey of object detection techniques in microorganism image analysis: from classical methods to deep learning approaches. Artif Intell Rev 56, 1627–1698 (2023). 10.1007/s10462-022-10209-1

21. Juan C. Miranda, Jordi Gené-Mola, Manuela Zude-Sasse, Nikos Tsoulias, Alexandre Escolà, Jaume Arnó, Joan R. Rosell-Polo, Ricardo Sanz-Cortiella, José A. Martínez-Casasnovas, Eduard Gregorio, (2023) Fruit sizing using AI: A review of methods and challenges, Postharvest Biology and Technology,Volume 206,2023,112587,ISSN 0925-5214, 10.1016/j.postharvbio.2023.112587.

22. Kim E, Hong SJ, Kim SY, Lee CH, Kim S, Kim HJ, Kim G. (2022) CNN-based object detection and growth estimation of plum fruit (Prunus mume) using RGB and depth imaging techniques. Sci Rep. 2;12(1):20796. doi: 10.1038/s41598-022-25260-9. PMID: 36460731; PMCID: PMC9718814.

23. K. Chotayapa, T. Leethamchayo, P. Chinnawong, T. Samernate, P. Nonejuie and T. Achakulvisut, “Deep Learning-Based Object Detection And Bacteria Morphological Feature Extraction For Antibiotic Mode Of Action Study,” 2023 15th Biomedical Engineering International Conference (BMEiCON), Tokyo, Japan, 2023, pp. 1-5, doi: 10.1109/BMEiCON60347.2023.10322010.

24. Stylianidou, S., Brennan, C., Nissen, S. B., Kuwada, N. J., & Wiggins, P. A. (2016). SuperSegger: robust image segmentation, analysis and lineage tracking of bacterial cells. Molecular microbiology, 102(4), 690–700. 10.1111/mmi.13486

25. Chen, Leiyu, Shaobo Li, Qiang Bai, Jing Yang, Sanlong Jiang, and Yanming Miao. (2021). “Review of Image Classification Algorithms Based on Convolutional Neural Networks” Remote Sensing 13, no. 22: 4712. 10.3390/rs13224712

26. Gulli, A., & Pal, S. (2017). Deep learning with keras: Implementing deep learning models and neural networks with the power of python. Packt Publishing.

27. Tan, M. & Le, Q.. (2019). EfficientNet: Rethinking Model Scaling for Convolutional Neural Networks. Proceedings of the 36th International Conference on Machine Learning, in Proceedings of Machine Learning Research 97:6105-6114 Available from https://proceedings.mlr.press/v97/tan19a.html.

28. Chen, Leiyu, Shaobo Li, Qiang Bai, Jing Yang, Sanlong Jiang, and Yanming Miao. (2021). “Review of Image Classification Algorithms Based on Convolutional Neural Networks” Remote Sensing 13, no. 22: 4712. 10.3390/rs13224712

29. Kingma, D. and Ba, J. (2015) Adam: A Method for Stochastic Optimization. Proceedings of the 3rd International Conference on Learning Representations (ICLR 2015).

30. S. Ren, K. He, R. Girshick and J. Sun. (2017) “Faster R-CNN: Towards Real-Time Object Detection with Region Proposal Networks” in IEEE Transactions on Pattern Analysis & Machine Intelligence, vol. 39, no. 06, pp. 1137–1149. doi: 10.1109/TPAMI.2016.2577031

31. Martín Abadi, Paul Barham, Jianmin Chen, Zhifeng Chen, Andy Davis, Jeffrey Dean, Matthieu Devin, Sanjay Ghemawat, Geoffrey Irving, Michael Isard, Manjunath Kudlur, Josh Levenberg, Rajat Monga, Sherry Moore, Derek G. Murray, Benoit Steiner, Paul Tucker, Vijay Vasudevan, Pete Warden, Martin Wicke, Yuan Yu, and Xiaoqiang Zheng. (2016). TensorFlow: a system for large-scale machine learning. In Proceedings of the 12th USENIX conference on Operating Systems Design and Implementation (OSDI’16). USENIX Association, USA, 265–283.

32. Kingma, D. and Ba, J. (2015) Adam: A Method for Stochastic Optimization. Proceedings of the 3rd International Conference on Learning Representations (ICLR 2015).

33. Canny, J. (1986). A computational approach to edge detection. IEEE Transactions on Pattern Analysis and Machine Intelligence, (6), 679–698.

34. Sionov, R. V., Banerjee, S., Bogomolov, S., Smoum, R., Mechoulam, R., & Steinberg, D. (2022). Targeting the Achilles’ Heel of Multidrug-Resistant *Staphylococcus aureus* by the Endocannabinoid Anandamide. International journal of molecular sciences, 23(14), 7798. 10.3390/ijms23147798

35. Anastasia S Kokoreva, Elena P Isakova, Vera M Tereshina, Olga I Klein, Natalya N Gessler, Yulia I Deryabina The Effect of Different Substrates on the Morphological Features and Polyols Production of Endomyces magnusii Yeast during Long-Lasting Cultivation Microorganisms. 2022 Aug 25;10(9):1709. doi: 10.3390/microorganisms10091709.

36. Haoming Liu, Hamid Karani, Jon Mallen, Weijie Chen, Arpan De, Sridhar Mani ^2^, Jay X Tang (Enterobacter sp. Strain SM1_HS2B Manifests Transient Elongation and Swimming Motility in Liquid Medium Zhiyu Zhang ^# 1^, Microbiol Spectr. 2022 Jun 29;10(3):e0207821. doi: 10.1128/spectrum.02078-21. Epub 2022 Jun 1.

37. Marilina Fernandez, Natalia S Paulucci, Eugenia Reynoso, Gustavo M Morales, Elizabeth Agostini, Paola S González Morphological and structural response of Bacillus sp. SFC 500–1E after Cr(VI) and phenol treatment. J Basic Microbiol. 2020 Aug;60(8):679-690. doi: 10.1002/jobm.202000076. Epub 2020 May 6.

38. Elizabeth A. Mueller, PLoS Genet. pH-dependent activation of cytokinesis modulates *Escherichia coli* cell size Conceptualization, Formal 2020 Mar; 16(3): e1008685. Published online 2020 Mar 23. doi: 10.1371/journal.pgen.1008685

